# Evolutionary rewiring of an ancient gene regulatory network specifies secretory cells in stony corals

**DOI:** 10.64898/2026.03.31.715602

**Authors:** Ryan M. Besemer, Laura Slama, Nicole Fogarty, Sarah Arnold, Koty Sharp, Leslie S. Babonis, Jacob F. Warner

**Affiliations:** Department of Biology and Marine Biology, University of North Carolina Wilmington; Wilmington NC, 28409, USA; University of North Carolina Wilmington Center for Marine Science; Wilmington NC, 28403, USA; Department of Ecology and Evolutionary Biology, Cornell University; Ithaca NY, 14853, USA; Department of Biology, Marine Biology, and Environmental Science, Roger Williams University; Bristol RI, 02809

## Abstract

Stony corals are the eponymous keystone organisms for the most diverse marine ecosystems on earth: coral reefs. Despite their importance, little is known about the developmental genetic programs that drive their embryonic development and subsequent emergence of coral-specific traits. Like all developmental processes, coral development is governed by a series of genetic interactions collectively termed a gene regulatory network. Evolutionary changes in these gene regulatory networks are key sources of biological innovation, but how these networks operate in corals and how they have changed over time remains poorly understood. Here, we integrate chromatin profiling and transcriptomics to define the cis-regulatory architecture of early development in stony corals using the emerging model, *Astrangia poculata*. We find that key elements of the deeply conserved cnidarian endomesoderm gene regulatory network have been co-opted into a novel subnetwork that specifies secretory cells in corals. Using transgenic reporter assays, we found that a proximal cis-regulatory element of the endomesodermal gene Brachyury is sufficient to drive coral-specific expression patterns in the sea anemone *Nematostella vectensis,* demonstrating these elements as a key evolvable unit in stony corals. Collectively these findings establish a gene-regulatory framework for coral development and cell specification and suggest that evolutionary rewiring of ancestral developmental programs is a key driver in the evolution and diversification of stony corals.

## Introduction

Much of the diversity of life is attributed to evolutionary changes in the gene regulatory networks that drive animal development. Comparing these networks among diverse organisms can provide a lens into how dGRNs can be re-wired to produce new cell types, forms, species, or life histories. The phylum Cnidaria, which includes jellyfish, sea anemones, hydroids, and corals, are an especially diverse group of organisms and are among the earliest branching metazoans, having diverged from bilaterians approximately 750 million years ago (McFadden et al., 2021a; Park et al., 2012). Most members of Anthozoa, a subphylum of Cnidaria that includes sea anemones and corals, develop via a free-swimming planula that eventually settles and metamorphoses into a sessile polyp. During the planula stage, anthozoan larvae express a conserved suite of developmental gene regulatory network (dGRN) genes with highly conserved expression patterns (Gilbert et al., 2024; Hayward et al., 2015a; Levy et al., 2021; Okubo et al., 2016a; Röttinger et al., 2012a; Schwaiger et al., 2022). In studies examining tropical coral larvae, ‘ectopic’ ectodermal expression of several endomesoderm dGRN genes, including FoxA, have been reported, although the significance of this expression has not been investigated (Hayward et al., 2015b; Okubo et al., 2016b). In this study, we provide evidence that this ectopic expression is part of a novel endomesodermal subnetwork specifying secretory cells in stony corals.

In this study we sought to identify the genetic programs driving the earliest stages of development of stony corals to understand the emergence of coral specific traits. To do this, we used the northern star coral, *Astrangia poculata,* as a model. This temperate coral is found in abundance in shallow waters off the east coast of the United States (Dimond et al., 2013). Recent phylogenomic analysis positions *Astrangia* among major tropical coral models (Fig. 1A; Vaga et al., 2025). *Astrangia* colonies are compact (Fig. 1B), and they can be induced to spawn during a wide reproductive season in the laboratory using a temperature shift, making *Astrangia* an ideal model system to study coral development (Ashey et al., 2025; Szmant-Froelich et al., 1980; Warner et al., 2025). Here, we systematically characterized gene expression and chromatin accessibility in developing *Astrangia* embryos to derive the regulatory landscape underlying germ layer specification. We show that during *A. poculata* development, several transcription factors canonically associated with endomesoderm specification exhibit derived ectodermal expression patterns where they are co-expressed with secretory genes. Perturbation of the endomesodermal network globally results in defects in endomesoderm formation in addition to down regulation of the secretory genetic program suggesting that specification of coral secretory cells depend on components of an ancestral endomesodermal network. Finally, we show that a cis-regulatory element from the conserved endomesodermal regulator Brachyury is sufficient to drive coral-like expression in the sea anemone *Nematostella vectensis*. Together our results suggest that during the evolution elements of the endomesoderm GRN were co-opted into a novel subnetwork to specify secretory cells.

**Figure 1:**
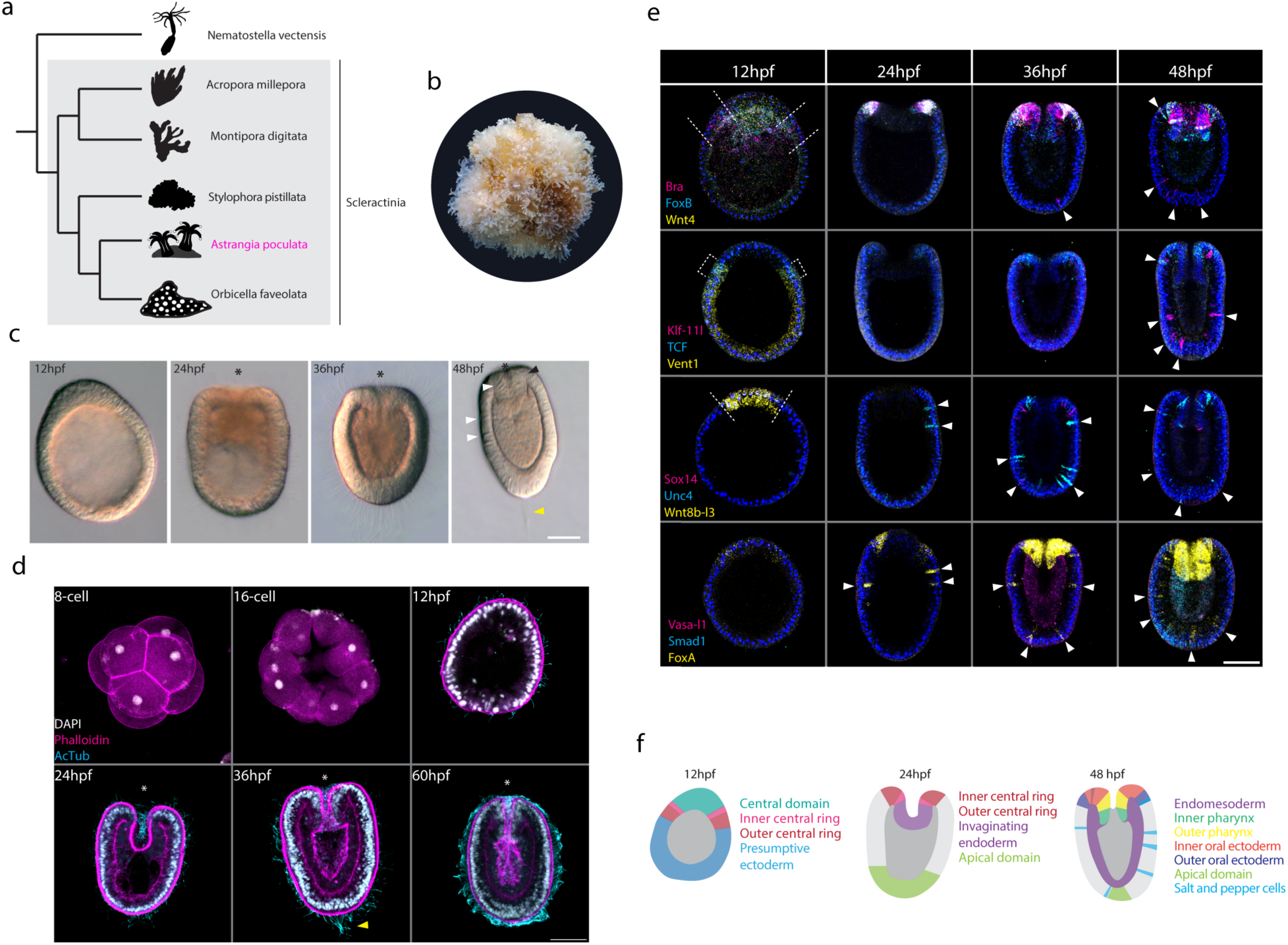
Highly conserved endomesoderm-associated genes are expressed in divergent patterns during development of the temperate coral Astrangia poculata. A. Phylogeny highlighting relationship of *A. poculata* to select stony corals (Scleractinia). B. An adult *Astrangia poculata* colony. Colony size is approximately 5 cm. C. DIC images of *A. poculata* development from 12 hpf through 48 hpf. White arrowheads denote the appearance of cnidocytes. D. Max confocal projections of developing *A. poculata* stained for nuclei (DAPI, white), actin (phalloidin, magenta), and cilia (anti-acetylated alpha tubulin antibody, cyan). Yellow arrowhead denotes the aboral apical tuft. Asterisks mark the site of gastrulation/oral pole. E. Expression of key genes during *A. poculata* development. Dashed lines demarcate early oral expression domains. White arrowheads denote ectodermal ‘salt and pepper’ patterns. F. Summary of expression domains in developing *A. poculata*.

## Results

### Conserved developmental network genes exhibit divergent expression patterns in early coral embryos

The northern star coral, *Astrangia poculata*, develops via a free swimming planktotrophic larva (Fig. 1C-D; Szmant-Froelich et al., 1980). After holoblastic cleavage, a ciliated swimming blastula forms at 12 hours post fertilization (12 hpf; Fig. 1C, D). Using *in situ* hybridization chain reaction (HCR) we found that the 12 hpf early blastula is broadly divided into two expression domains marking the future oral and aboral regions. The presumptive oral pole is further divided into concentric nested domains that express Wnt8b-like 3 (central domain), Wnt4 (central domain), FoxB (inner central ring), Brachyury (outer central ring), and TCF (outer central ring; Fig. 1E, F). The transcription factor Vent1 is broadly expressed in the aboral region and overlaps with the outer central ring (Fig. E). A large number of transcripts corresponding to Wnt signaling pathway members, including Beta-catenin, Axin, Flamingo, and several Wnt ligands, are present at high levels, suggesting that similar to what has been shown in the sea anemone, *Nematostella vectensis,* canonical Wnt signaling is a key player in germ layer specification (Supp. Fig. 2; Haillot et al., 2025; Leclère et al., 2016; Marlow et al., 2013; Röttinger et al., 2012).

**Figure 2:**
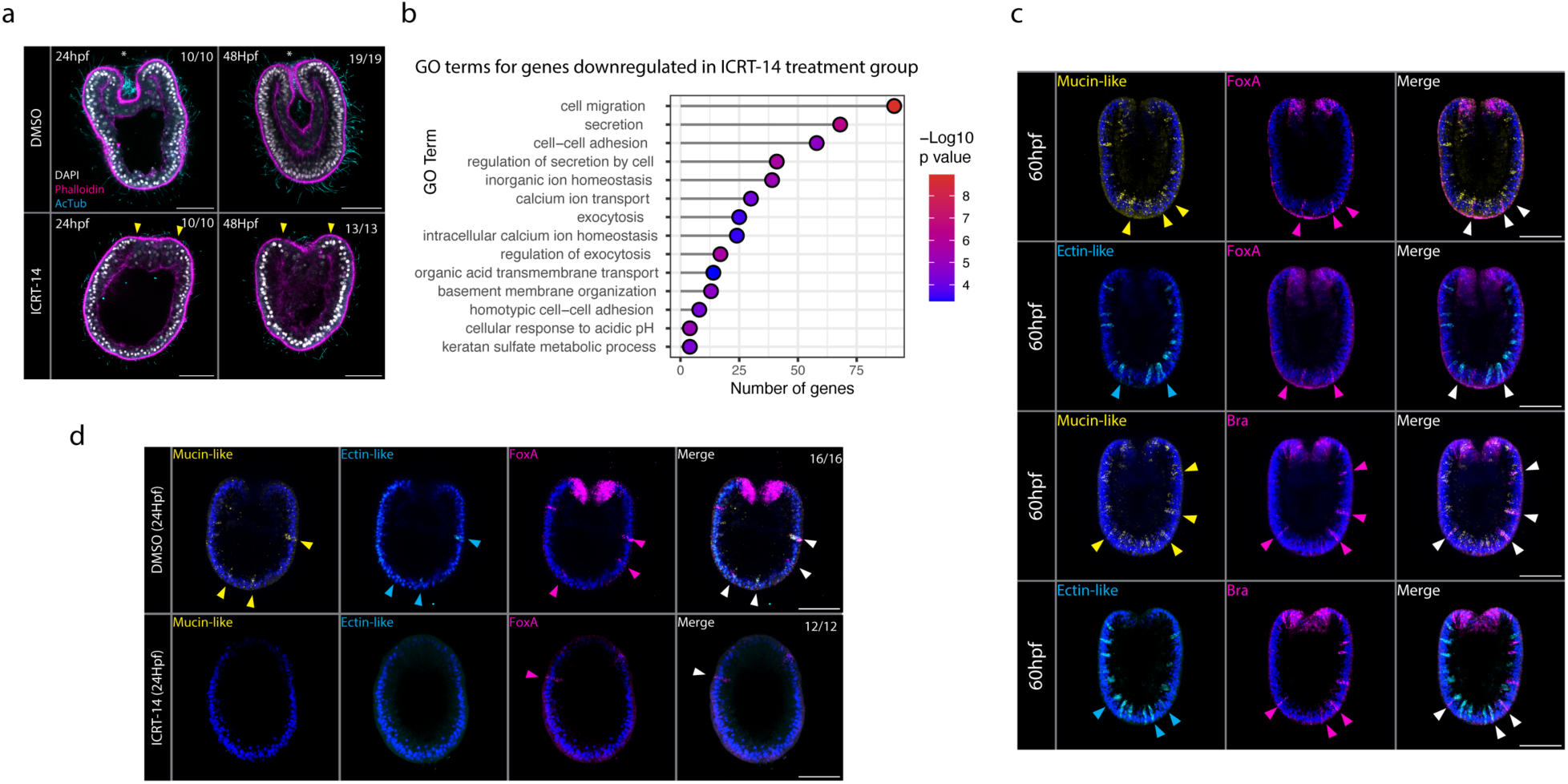
Blocking endomesoderm formation prevents secretory cell specification. A. *Astrangia poculata* treated with the TCF inhibitor ICRT-14 compared to DMSO controls stained for nuclei (DAPI, white), actin (phalloidin, magenta), and cilia (anti-acetylated alpha tubulin antibody, cyan). Asterisks mark the site of gastrulation/oral pole. Yellow arrowheads denote thickening of the oral pole but lack of gastrulation. B. GO terms linked to secretory processes are significantly downregulated in ICRT-14 treatment group versus DMSO control. C. FoxA and Brachyury are co-expressed with the secretory genes Mucin-like and Ectin-like. D. Expression of FoxA, Ectin-like, and Mucin-like are downregulated in ICRT-14 treated embryos. Arrowheads denote ectodermal expression domains.

At 24 hpf, the embryo gastrulates and the presumptive endomesoderm begins to invaginate (Fig. 1 C, D). Several genes are expressed at the site of invagination including FoxB, Brachyury, Wnt4, and TCF. By 36hpf, gastrulation is complete, and the larval pharynx begins to form. Similar to what has been observed in *Nematostella vectensis*, the *Astrangia* pharynx expresses FoxA, and Brachyury is expressed at the margin of the invaginating pharynx where the tissue transitions to oral ectoderm (Fig. 1E; Fritzenwanker et al., 2004; Haillot et al., 2024; Martindale et al., 2004; Schwaiger et al., 2022; Steinmetz et al., 2017). Interestingly, in sharp contrast to what has been observed in *Nematostella*, several genes associated with endomesoderm formation, including Brachyury, Klf1, FoxA, and TCF, are expressed in scattered cells throughout the aboral ectoderm (Fig. E; Marlow et al., 2013; Röttinger et al., 2012b), suggesting these genes participate in a novel network arrangement to specify additional cell types in corals.

### Blocking the endomesoderm network results in downregulation of secretory genes

In the sea anemone *Nematostella vectensis*, β-Catenin/TCF mediated Wnt signaling is critical to activate endomesoderm-associated genes, and inhibiting TCF leads to downregulation of the endomesodermal network genes, FoxA and Brachyury (Haillot et al., 2025; Marlow et al., 2013; Röttinger et al., 2012a; Wijesena et al., 2017). We, therefore, sought to investigate if manipulating this pathway had an equivalent effect in *Astrangia,* modulating expression of endomesodermal target genes in both oral regions and ectopic ectodermal domains. To do this, we treated fertilized *Astrangia poculata* zygotes with ICRT-14, a potent inhibitor of β-Catenin/TCF mediated signal transduction in vertebrates and cnidarians (Gonsalves et al., 2011; Marlow et al., 2013). Embryos treated with 20 μM ICRT-14 from fertilization failed to gastrulate (Fig. 2A). Using RNAseq, we observed a significant downregulation of genes with gene ontology (GO) mappings related to secretion in the ICRT-14 treated group (Fig. 2B). Given this, we examined the expression of the two genes associated with coral secretory cells, Ectin-like, a CAP-domain-containing secreted protein, and Mucin-like, an extracellular matrix protein containing several Thrombospondin type 1 repeats, both of which are expressed in secretory cells in the larvae of the tropical coral *Stylophora pistillata* (Levy et al., 2021). Using HCR, we examined the expression of Mucin-like and Ectin-like and found these genes to be expressed in a salt and pepper pattern in the aboral ectoderm and in partial overlap with the ectodermal expression domain of Brachyury and FoxA (60hpf; Fig. 2C), suggesting that Brachyury and FoxA positive cells in the ectoderm are likely secretory cells. Embryos treated with 20 μM ICRT-14 from fertilization exhibited significant downregulation of FoxA in the oral and ectodermal domains in addition to abolished expression of the secretory genes Ectin-like and Mucin-like (Fig. 2D). These results suggest a dual role of the endomesodermal GRN in establishing germ layer identity and specifying ectodermal secretory cells.

### The regulatory landscape surrounding developmental network genes is highly conserved among corals

Given the striking gain of a novel expression pattern for endomesodermal genes in *Astrangia* embryos, we sought to better understand the regulatory mechanisms driving expression of these genes. To do this, we performed the Assay for Transposase-Accessible Chromatin with high-throughput sequencing (ATAC-seq) to identify open chromatin regions (OCRs) representing putative *cis*-regulatory elements (CREs) during embryogenesis of *A. poculata.* We performed ATAC-seq at the blastula stage (12 hpf), gastrula stage (24 hpf), and early planula stage (36 hpf). Our analysis revealed a dramatic increase in genome accessibility between 12 hpf and 24 hpf (Fig. 3A-D), consistent with activation of the zygotic genome. The largest fraction of these accessible regions (44%) is found within putative promoters (+/− 2 kb of the transcription start site), compared to 25% in intergenic regions, 18% in introns, and 8% in exons (Fig. 3A). This is similar to patterns observed in other cnidarians (Cazet et al., 2023; Sebé-Pedrós et al., 2018; Schwaiger et al., 2014). Finally annotated each peak for putative transcription factor binding sites by scanning the genome for known and novel transcription factor binding motifs (herein termed TFBs; van Heeringen & Veenstra, 2011).

**Figure 3:**
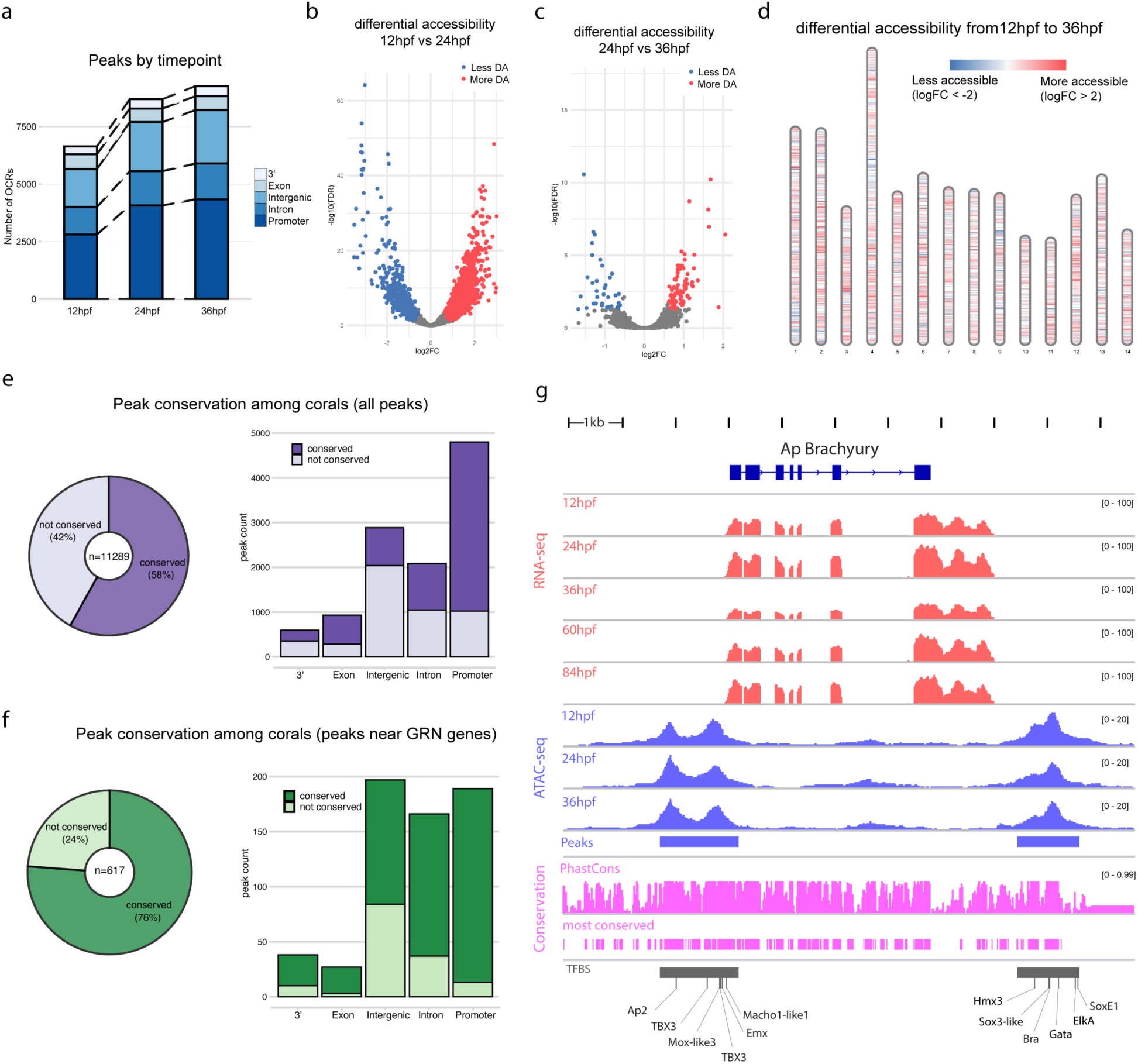
The cis-regulatory landscape of stony corals is highly conserved. A. ATAC-seq peaks by timepoint. B-D. differential peak accessibility analysis shows global increase in chromatin accessibility during development. E. ATACseq peak sequences are largely conserved among stony corals. F. Peaks near dGRN genes are highly conserved. G. The regulatory landscape surrounding the *Astrangia poculata* Brachyury locus with RNAseq coverage (red) and ATACseq coverage (periwinkle), and conservation analysis (pink).

We next examined the level of sequence conservation under each ATAC-seq peak. To do this we performed a whole genome alignment of the *A. poculata genome* with five tropical coral species chosen for their high-contiguity chromosomal-level genome assemblies (*Montipora capitata*, *Acropora millepora*, *Stylophora pistillata*, *Galaxea fascicularis*, and *Orbicella franski*). We then used PhastCons to score ATAC-seq peaks for conservation (defined as peaks overlapping with a PhastConst ‘most conserved’ element). We found that *A. poculata* ATAC-seq peak sequences were generally well conserved (Fig. 3E; 58%, n=11289) with those falling in promoter regions and exons exhibited the highest degree of conservation. ATAC peaks near GRN genes exhibited a much higher degree of conservation (Fig. 3F; 76%, n=617) with those in the immediate gene regions (promoter, intronic, exonic, and 3’ peaks) exhibiting high rates of conservation compared to those in intergenic regions.

### Proximal cis-regulatory elements are a key evolvable unit in corals

As changes in *cis-*regulatory elements are key drivers of biological innovation and evolution (Davidson et al., 2022; Devens et al., 2023; Uebbing et al., 2024; Wittkopp & Kalay, 2012), we sought to identify the regulatory elements responsible for the novel expression of endomesodermal genes in developing *Astrangia.* For this, we focused on the *cis-*regulatory program of the *Astrangia poculata* Brachyury locus. Brachyury is a critical node in the cnidarian embryonic gene regulatory network and regulates morphogenesis of the invaginating pharynx in corals and sea anemone embryos (Hayward et al., 2015a; Okubo et al., 2016a; Servetnick et al., 2017a; Yasuoka et al., 2016). In *A. poculata*, Brachyury is expressed at the margin of invaginating endomesoderm during gastrulation (Fig. 1E). Later, beginning at 24 hpf, it is expressed in the pharynx and in scattered cells throughout the aboral ectoderm (Fig. 1E, 4A). This pattern is in sharp contrast to what has been observed in the starlet sea anemone, where Brachyury is restricted to the developing oral pole and has no detectable expression in the ectoderm (Fig. 4A, 4B; Martindale et al., 2004, Fritzenwanker et al., 2004).

**Figure 4:**
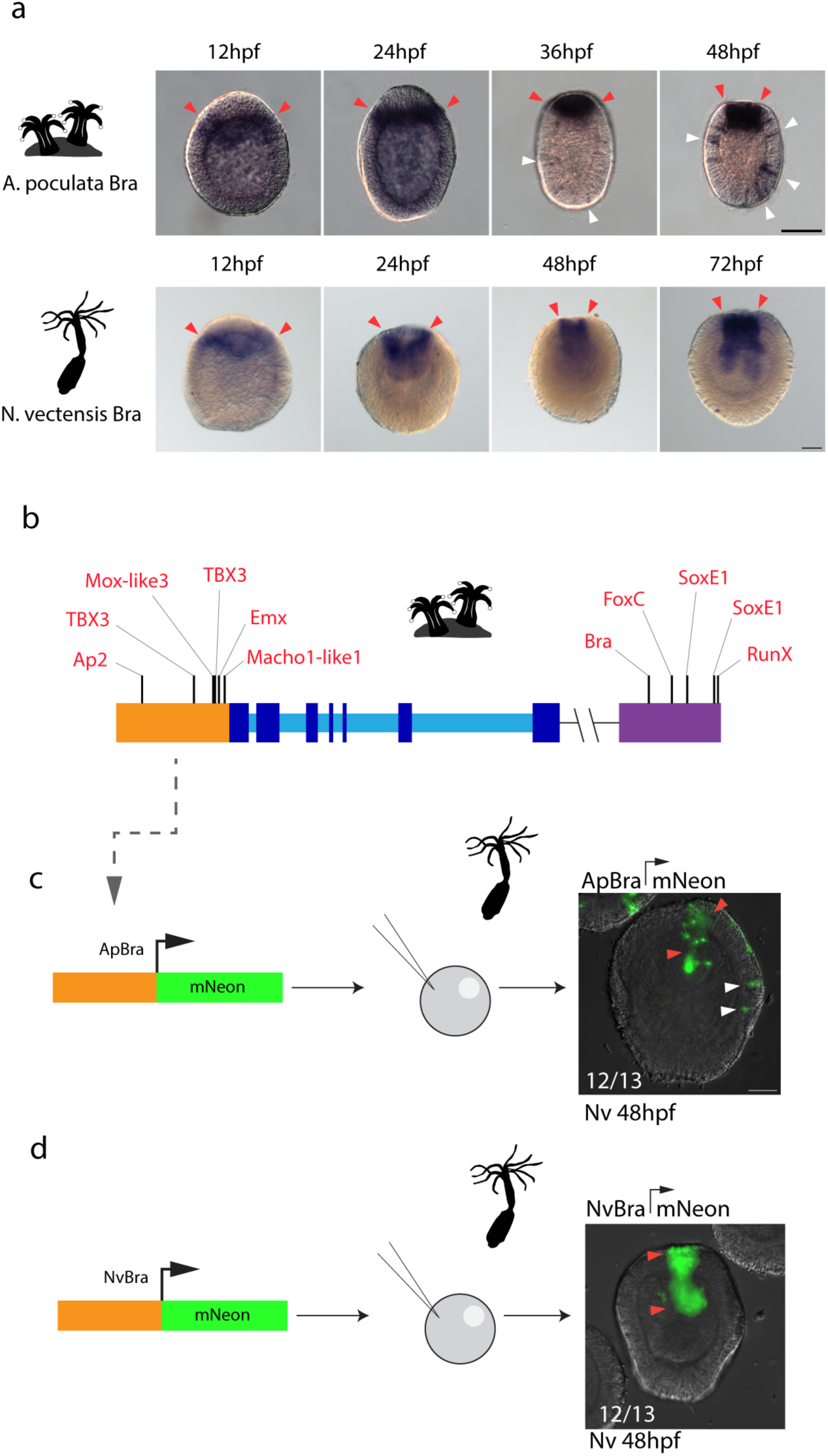
A 5’ cis-regulatory module of the Astrangia Brachyury promoter is sufficient to drive ectodermal expression in the sea anemone, Nematostella vectensis. A. Comparison of Brachyury expression in developing *Astrangia poculata* and the sea anemone, *Nematostella vectensis*. Red arrowheads denote oral expression domains; white arrowheads highlight ectodermal domains. B. Summary of the *A. poculata* gene locus with transcription factor binding sites annotated in the 5’ (orange) and 3’ (purple) regulatory modules. C. Summary of cross-species expression assay: The 5’ regulatory module from the *A. poculata* gene was cloned in-frame with mNeon into a meganuclease transgenesis plasmid and injected into fertilized *N. vectensis* eggs. Resultant chimeric F0 embryos exhibited oral (red arrowheads) and ectodermal (white arrowheads) expression domains. D. A 2 kb region 5’ of the of *N. vectensis* Brachyury promoter cloned in-frame with mNeon into the pNvt meganuclease transgenesis vector and injected into fertilized *N. vectensis* eggs. Fluorescence was assayed at 48 hpf.

We examined in detail the *cis*-regulatory landscape surrounding Brachyury in *A. poculata.* We found an open chromatin region spanning a 2 kb region of the proximal promoter and a putative regulatory element approximately 2 kb downstream of the coding region (Fig. 3G). As cnidarian regulatory information is frequently encoded in the proximal gene promoter (Cazet et al., 2023; Sebé-Pedrós et al., 2018; Schwaiger et al., 2014), we sought to test whether the cis-regulatory element found in the *A. poculata* Brachyury promoter could drive the Brachyury expression pattern in the sea anemone *N. vectensis* (estimated to have diverged from the *A. poculata* lineage approximately 650 million years ago; McFadden et al., 2021b; Park et al., 2012). To do this, we assembled a series of transgenic reporter constructs consisting of the *A. poculata* Brachyury regulatory regions driving expression of the fluorescent protein, mNeon, flanked by iSceI meganuclease sites. When we injected this construct into fertilized eggs of *N. vectensis* the we observed that this construct recapitulated the *A. poculata* Brachyury expression pattern in the pharynx (Fig. 4C, white arrows) and exhibited ectopic expression in the ectoderm (Fig. 4C, red arrows). By contrast, injecting a construct carrying a 2 kb fragment of the *N. vectensis* Brachyury promoter upstream of a fluorescent protein recapitulated the endogenous *N. vectensis* expression in the oral and pharyngeal regions only (Fig. 4D). Collectively, these results identify the proximal promoter of Brachyury as the evolvable unit responsible for the ectopic expression of this gene in *Astrangia* as it became co-opted into an endomesodermal subnetwork specifying secretory cells.

## Discussion

### Key endomesodermal GRN nodes have been re-wired along the coral lineage to specify secretory cells

This study points to a model in which key endomesodermal dGRN genes have been re-wired to specify secretory cells in developing scleractinian coral embryos. In the sea anemone *Nematostella,* FoxA, TCF, and Brachyury are exclusively expressed in oral regions: FoxA and TCF are expressed in the blastopore and presumptive endoderm, while Brachyury is expressed in the invaginating pharynx (Fritzenwanker et al., 2004; Martindale et al., 2004). In contrast, we found that in the temperate coral, *Astrangia poculata,* these genes are expressed in developing endomesoderm in addition to a secondary expression domain of scattered ectodermal cells that secretory genes, including Mucin-like and Ectin-like (Fig. 1E, 2C, 4A). Furthermore, we found that globally inhibiting the endomesodermal GRN by knocking down TCF results in downregulation of genes involved in secretion (Fig. 2C-E), consistent with a novel role for the endomesodermal GRN in specifying secretory cells in stony corals.

This study is not the first to report endomesoderm-associated genes being expressed in the ectoderm of developing coral embryos. In the tropical coral species, *Acroropora millepora*, several endomesoderm dGRN genes, including FoxA, exhibit ectodermal expression in addition to ‘canonical’ expression in the developing endomesoderm (Hayward et al., 2015b; Okubo et al., 2016b). A study examining expression of Brachyury in *Acropora millepora*, on the other hand, did not report ectodermal expression (Yasuoka et al., 2016). It is unclear if this is due to true differences in expression or due to the limitations of gene expression assays in *Acropora* embryos which have a high lipid content. Additional evidence linking the endomesoderm-associated genes to secretory cell specification in tropical corals comes from Levy et al., who performed single cell RNA-sequencing at larval and settled polyp stages in the smooth cauliflower coral, *Stylophora pistillata* (Levy et al., 2021). While the authors of that study did not discuss endomesoderm-associated genes specifically, searching their dataset identified Brachyury as enriched in secretory gland cells of larvae and FoxA as enriched in calicoblasts (skeleton secreting cells) of settled polyps (Supp. Fig. 3). Together these studies suggest that the co-option of endomesodermal network components to specify secretory cells is a widespread phenomenon among scleractinian corals.

The divergent expression patterns of endomesoderm-associated genes in *Astrangia* indicate that the gene regulatory landscape of the endomesodermal GRN is not immutable and has been re-wired during evolution of stony corals. Surprisingly, this rewired regulatory logic can be interpreted by the sea anemone *Nematostella vectensis*, which diverged from the *A. poculata* lineage approximately 650 million years ago (Fig. 3C). These results point to the *cis*-regulatory architecture of developmental GRN genes as a key evolvable unit among Anthozoa and demonstrate how a dGRN can be rewired to drive diversification of stony corals.

## Acknowledgements

This work was supported by the North Carolina Biotechnology Center (2022-FLG-3803, J.F.W.), the National Institutes of Health (R15GM139113-01A1 to J.F.W. and R35GM147253-01 to L.S.B.), and the Coral Research and Development Accelerator Platform (CORDAP, J.F.W.) and a Marine Biological Laboratory Whitman fellowship to J.F.W. K.S. was supported by an Early Career Development Award to K.S. (Institutional Development Award (iDeA) Network for Biomedical Research Excellence, NIGMS, in the National Institutes of Health (P20GM103430)).

## Author contributions

JFW and RMB conceived the study. JFW, RMB, LS, SA, and LSB performed experiments. JFW and RMB analyzed data. NF and KS provided animal colonies and animal expertise. JFA, LSB, KS, and NF provided funding. JFW and RMB wrote the manuscript with input from all authors. All authors reviewed and approved the manuscript.

## Competing interests

Authors declare that they have no competing interests.

### Data, code, and materials availability

Raw ATACseq and RNAseq reads are available at the NCBI Short Read Archive (PRJNA1456634). Data files from the analysis are included in the supplementary materials. Code for generating the figures is available at https://github.com/warnerlab/Besemer_et_al_2026

## Supplementary Materials

### Materials and Methods

#### Animal collection spawning and larval rearing

*Astrangia poculata* colonies were collected at Fort Wetherhill, Rhode Island, USA and transferred to Roger Williams University. To induce spawning, the colonies were heat shocked by transferring them to 26°C for 1 hour. Colonies then spawned approximately 90 minutes later. Eggs and sperm were collected separately and combined in a small volume for fertilization. Fertilized eggs were then transferred to 30ml petri dishes for larval culture.

*Nematostella vectensis* were provided by Sarah Tulin (Canisius College) and maintained in closed culture in the laboratory. Animals were spawned as previously described (Genikhovich & Technau, 2009).

#### Antibody staining

Embryos were fixed in 4% Paraformaldehyde in PBS for 1 hour at room temperature. The embryos were then washed four times in PBSTween 0.1% for 10 minutes. Embryos were then blocked for 1 hour in 10% Normal Goat Serum (NGS) and incubated in 10% NGS containing anti-acetyl-alpha tubulin (Invitrogen 322700) at 1:200 overnight at 4°C. Next, the embryos were washed four times in PBSTween 0.1% for 10 minutes and then incubated in 10% NGS containing Alexa Fluor 488 conjugated goat anti mouse secondary antibody (AbCam ab150113) at 1:200 for four hours are RT. Embryos were washed four times in PBSTween 0.1% for 10 minutes and were then counterstained by incubating in a 10% NGS solution containing Alexa Fluor 546 conjugated Phalloidin (Invitrogen A22283) at 1:200 and DAPI 1:2500 overnight at 4°C. Embryos were washed four times in PBSTween 0.1% for 10 minutes and mounted in 80% glycerol for imaging.

#### Pharmacological treatments

100 fertilized single cell zygotes were transferred to 5ml ASW containing either 0.1% DMSO (control) or 0.1% DMSO containing iCRT-14 (MCE HY-16665) at a final concentration of 20mM. This concentration we determined by testing a range of doses (5mM, 10mM, 20mM, 40mM) for a concentration that reliably produced a phenotype without producing toxicity. To avoid attributing a phenotype to toxicological developmental delay, embryos were imaged at 24hpf and 48hpf. Each treatment was performed in triplicate.

#### In situ hybridization and in situ hybridization chain reaction

Gene expression was localized using *in-situ* hybridization (ISH). Embryos at desired developmental stages were first fixed in 4% PFA (paraformaldehyde) overnight at 4°C. A series of six washes was performed on the embryos in PBSTr 0.2% for tropical embryos (phosphate buffered saline with TritonX 100) or PBSTw 0.1% for temperate embryos (phosphate buffered saline with Tween-20). The embryos are then transferred to ice cold methanol and stored at -20°C.

*A. poculata* embryos were first stepped out of methanol into TBST. After one full wash in TBST for five minutes, embryos were allowed to prehybridize in hybridization buffer with 5% dextran sulfate at 65°C for at least one hour. While this occurred, the probe was prepared to a concentration of 0.5ng/ul at 65°C. The prehybridization buffer was removed and replaced with the hybridization and probe mix, allowing the embryos to hybridize overnight at 65°C. Following hybridization, the embryos were washed out of hybridization buffer first with two 15-minute washes of hybridization buffer without 5% dextran sulfate at 65°C. The embryos were then stepped out of hybridization buffer and into SSCT with a 50% hybridization/50% 2X SSCT wash, a 2X SSCT wash, a 0.2X SSCT wash, and a 0.1X SSCT wash all at 65°C. The embryos were finally washed in TBST for five minutes at room temperature. The TBST wash was followed with allowing the embryos to block in blocking solution (0.5% BSA and 2% heat inactivated sheep serum in TBST) for at least an hour. The embryos were then incubated overnight with the Anti-DIG antibody from Roche diluted to 1:2000 overnight at 4°C. The next day, embryos were washed with TBST twice for fifteen minutes, followed by an additional four washes for ten minutes all at room temperature. The embryos were then washed with an alkaline phosphatase buffer twice before incubating in alkaline phosphatase buffer with NBT (6.75ul/ml) and BCIP (3.5ul/ml). The embryos were monitored during the NBT/BCIP development every thirty minutes with fresh solution every hour for the first day. Development was allowed to proceed until a signal was noticed in the embryos. If the gene took longer than a day to develop, the embryos were placed back into alkaline phosphatase buffer without NBT or BCIP at 4°C overnight. To conclude development, embryos were washed five times in PBSTw 0.1% at room temperature. Samples were then mounted on slides and imaged using a Leica Thunder Imager.

In addition to traditional ISH, the hybridization chain reaction (HCR) was utilized to localize gene expression. HCR probes were designed and ordered from Integrated DNA Technologies (IDT). Embryos were fixed in 4% PFA at 4°C overnight, following the same fixative process as those used in traditional ISH. Just as before, embryos were washed a total of six times in PBSTw 0.1% before being stored at -20°C in methanol.

Embryos were then retrieved from -20°C and placed in a well plate for HCR washes. First, embryos were stepped out of methanol and into PBSTw 0.1%. Following a complete wash in PBSTw 0.1%, embryos were washed an additional five times in PBSTw 0.1% for five minutes at room temperature. Meanwhile, the hybridization buffer from Molecular Instruments was thawed to 37°C. Following the PBSTw 0.1% washes, the embryos were washed three times in 5X SSCT for five minutes at room temperature. Embryos were pre-hybridized for at least an hour in 50ul of probe hybridization buffer at 37°C. Meanwhile the probe was prepared by adding 0.4pmol to 50ul of probe hybridization buffer. After pre-hybridization, the probe and probe hybridization mix was added bringing the total volume to 100ul. The embryos were allowed to hybridize at 37°C overnight. The next day, probe wash buffer was added to each well for fifteen minutes at 37°C. The probe was washed out with probe wash buffer from molecular instruments using two twenty-minute washes and two thirty-minute washes all at 37°C. Two 5X SSCT washes were performed on the embryos at room temperature. During the SSCT washes, the hairpins were prepared by heat shocking them to 95°C for 90 seconds and allowing them to cool to room temperature in the dark. Right before their addition, the hairpins were added to 100ul of amplification buffer from molecular instruments at room temperature. As much SSCT as possible was removed and the embryos were placed in the hairpin mixture overnight in the dark at room temperature. The next day, 5X SSCT was added to each well and allowed to incubate for ten minutes at room temperature. Excess hairpins were then removed with two ten-minute washes and two thirty-minute washes in 5X SSCT at room temperature. The embryos were then washed five additional times at room temperature in 5X SSCT for ten minutes before suspending embryos in 30% glycerol/70% 5X SSCT with DAPI at 1:5000. Slides were then prepared and samples were imaged on a Leica SP8 Confocal microscope.

#### Ortholog identification and GRN gene model curation

*Astrangia poculata* gene orthologs were assigned using Orthofinder (Emms & Kelly, 2019) and the predicted protein sequences from the *A. poculata* genome assembly (Stankiewicz et al., 2025), the *Orbicella faveolata* genome assembly (RefSeq GCF_002042975.1), and the *Nematostella vectensis* genome assembly (Zimmermann et al., 2023). A list of 276 dGRN genes was curated using the Orthofinder results, and reciprocal best blast hit of *A. poculata* proteins and known developmental genes previously identified in *Nematostella vectensis*.

#### RNAseq analysis

RNA was collected by pelleting embryos (300 embryos for *Astrangia,*150 embryos for *Orbicella*) removing sea water and adding 500ml of Tri reagent. Samples were immediately homogenized with a pestle and snap frozen in liquid nitrogen for storage at -80C. Total RNA was purified using the Direct-zol RNA Miniprep Kit (Zymo Research R2050) prior to being sent to Novogene for Illumina sequencing. For each sample, 75 base pair paired end sequencing was carried out on the Illumina platform. Resultant reads were aligned to the *Astrangia poculata* genome (Stankiewicz et al., 2025) or the Orbicella faveolata genome (NCBI GCF_002042975.1) using Hisat2 (version 2.1.0) and quantified using featureCounts (version 1.6.0).

#### ATACseq library preparation and sequencing

ATAC-seq was performed following the Omni-ATAC-seq protocol (Grandi et al., 2022). Each sample was lysed and approximately 50,000 nuclei were isolated for the transposition reaction using the Illumina TDE1 enzyme and buffer (catalog number 20034197). Sequencing libraries were amplified using qPCR and the NEBnext enzyme kit (catalog number E7645S). Libraries were purified using a Qiagen Minelute kit (catalog number 28004) and size selected using AmpureXP beads (Beckman Coulter) at a 1.8:1 ratio of beads to library. Libraries were then pooled and sequenced on an Illumina NextSeq 500 machine with 75BP-PE reads.

#### ATACseq data analysis

Reads were filtered for illumine adapters using skewer (version 0.2.2) and aligned to the *Astrangia poculata* genome (Stankiewicz et al., 2025) using bwa-mem (version 0.7.17). Reads mapping to mitochondrial sequences were removed using samtools and filtered for mapping quality and maximum fragment length of 850Bp using alignmentSieve from the deepTools package (version 3.5.6). Coverage tracks were generated using bamCoverage from the deepTools package (version 3.5.6). Peaks were called using Genrich (version 0.6.1). ATAC peaks were annotated using annotatePeaks.pl from the HOMER package (version 4.11). Peak quantification was performed using featureCounts (version 1.6.0). Differential peak analysis was performed using edgeR (version 3.19) using reads mapped to the TSS regions to calculate the normalization factors. The resultant fold change was used as input into RIdeogram (version 0.2.2) to generate chromosome plots. Transcription factor binding sites (TFBS) were annotated using gimme scan and gimme motifs2factors from the GimmeMotifs tools suite (version 0.18.0).

#### Reporter assays

A 2kb region upstream of the transcriptional start site of *Astrangia poculata* Brachyury or *Nematostella vectensis* Brachyury was synthesized as a geneblock by IDT and cloned into a pNVT meganuclease transgenesis vector using AscI and PacI restriction sites to clone the fragment in front of the mNeon sequence. Purified plasmid was injected into fertilized *N. vectensis* eggs at 25ng/μl along with rhodamine dextran dye, as described previously (Renfer & Technau, 2017). Resultant embryos were then fixed at 48hours post fertilization in 4% PFA for 20 minutes at room temperature. The fixative was washed 2X in PBS before imaging.

#### Bioinformatic programs used

| <u>Task</u> | <u>Program</u> | <u>version</u> |
| --- | --- | --- |
| RNAseq read alignment | Hisat2 | 2.1.0 |
| RNAseq quantification | featureCounts | 1.6.0 |
| Coverage track generation<br>(bamCoverage) | deepTools | 3.5.6 |
| ATAC alignment filtering<br>(alignmentSieve) | deepTools | 3.5.6 |
| Filter ATACseq reads | Skewer | 0.2.2 |
| ATAC peak quantification | featureCounts | 1.6.0 |
| ATAC read mapping | Bwa-mem | 0.7.17 |
| ATAC peak calling | Genrich | 0.6.1 |
| BAM file processing | samtools | 1.3 |
| ATAC peak annotation<br>(annotatePeaks.pl) | HOMER | 4.11 |
| ATAC peak differential<br>analysis | R package: edgeR | 3.19 |
| TFBS annotation | GimmeMotifs | 0.18.0 |
| Chromosome painting | R package: RIdeogram | 0.2.2 |
| Genome masking | windowmasker | 2.2.22 |
| Ortholog detection | OrthoFinder | 2.5.5 |
| Tandem repeat<br>identification | trf | 4.09.01 |
| Various manipulation | bedtools | 2.28.0 |
| Multi-genome alignment | Cactus | 9.1.2 |
| Genome conservation<br>scoring | phastCons | 1.6 |

## Supplementary figures

**Supplemental figure 1:**
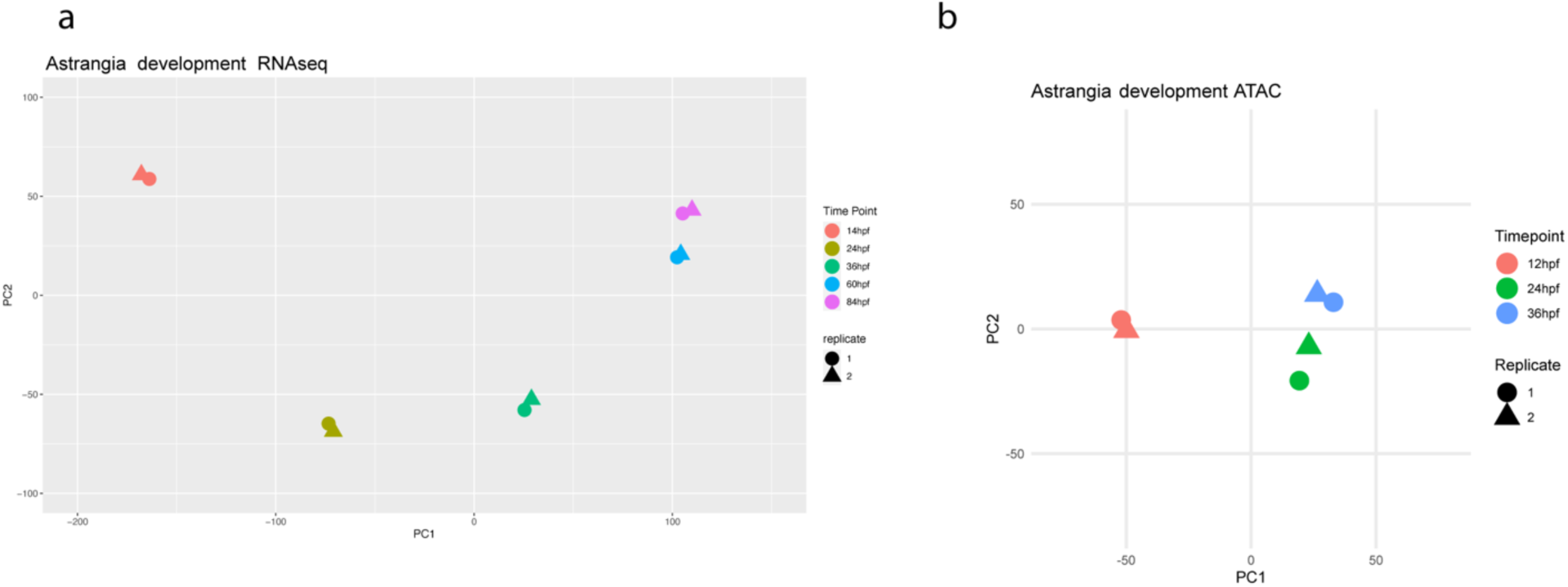
Principal component analysis of RNAseq (A) and ATACseq (B) of developing *A. poculata* embryos.

**Supplemental figure 2:**
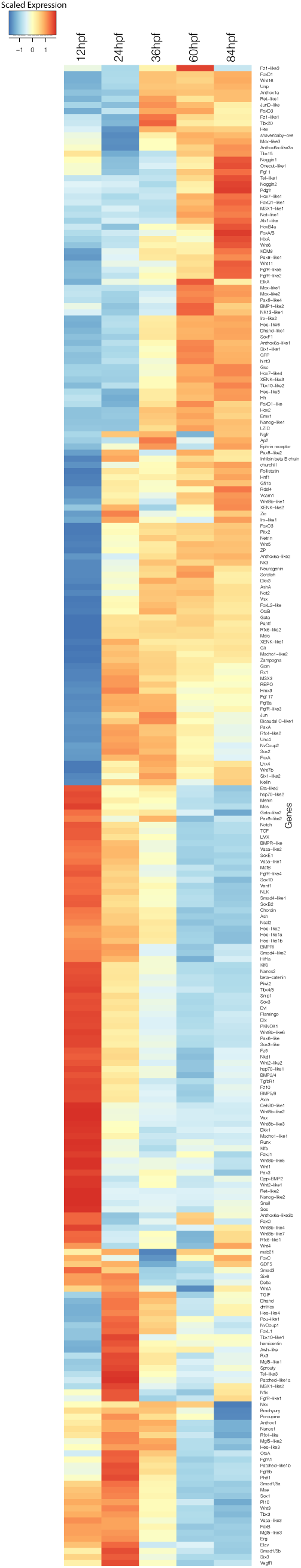
Heatmap of RNAseq results of curated dGRN genes.

**Supplemental figure 3:**
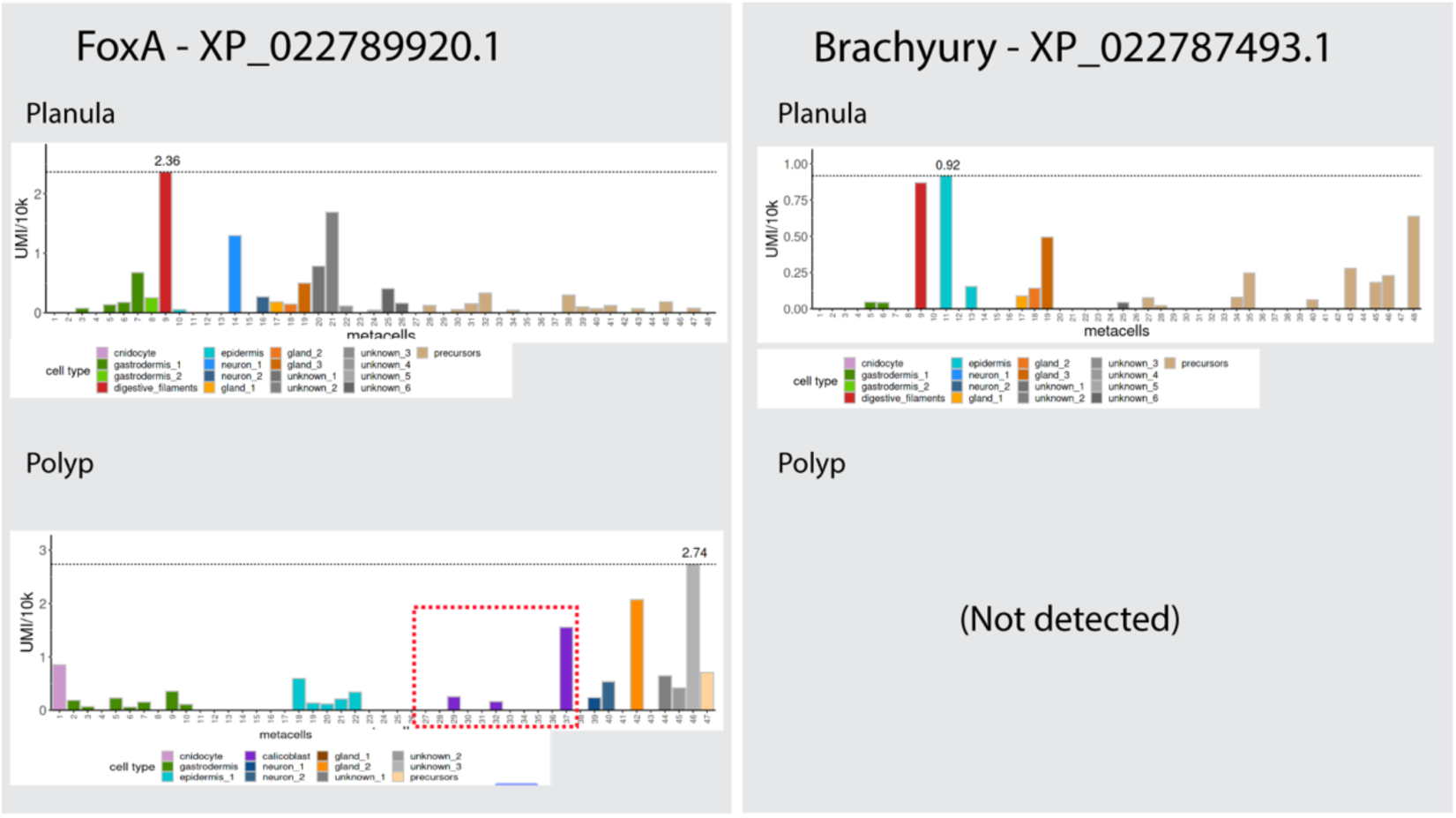
*Single cell RNAseq expression of FoxA, and Brachyury in the tropical coral Stylophora pistillata. Stylophora pistillata* orthologs were identified by reciprocal best blast between *A. poculata* and *S. pistillata*. Single cell RNAseq results from Levvy et al. 2021 were accessed, plotted and downloaded via the web app created by the original study authors found at: https://sebe-lab.shinyapps.io/Stylophora_cell_atlas/ (last accessed February 4^th^ 2026). Data S1. GO terms for genes significantly downregulated in ICRT treatment. Signficance threshold is logFC < -1.5 and FDR 0.05 in DMSO versus ICRT treatment. https://github.com/warnerlab/Besemer_et_al_2026/blob/main/RNAseq/GO_clust_ICRT_ TOpGO_fisher_BP.txt

